# Loss of function in the autism and learning disabilities associated gene *Nf1* disrupts corticocortical and corticostriatal functional connectivity in human and mouse

**DOI:** 10.1101/618223

**Authors:** Ben Shofty, Eyal Bergmann, Gil Zur, Jad Asleh, Noam Bosak, Alexandra Kavushansky, F. Xavier Castellanos, Liat Ben-Sira, Roger J. Packer, Gilbert L. Vezina, Shlomi Constantini, Maria T. Acosta, Itamar Kahn

## Abstract

Children with the autosomal dominant single gene disorder, neurofibromatosis type 1 (NF1), display multiple structural and functional changes in the central nervous system, resulting in neuropsychological cognitive abnormalities. Here we assessed the pathological functional organization that may underlie the behavioral impairments in NF1 using resting-state functional connectivity MRI. Coherent spontaneous fluctuations in the fMRI signal across the entire brain were used to interrogate the pattern of functional organization of corticocortical and corticostriatal networks in both NF1 pediatric patients and mice with a heterozygous mutation in the *Nf1* gene (*Nf1*^+/-^). Children with NF1 demonstrated abnormal organization of cortical association networks and altered posterior-anterior functional connectivity in the default network. Examining the contribution of the striatum revealed that corticostriatal functional connectivity was altered. NF1 children demonstrated reduced functional connectivity between striatum and the frontoparietal network and increased striatal functional connectivity with the limbic network. Awake passive mouse functional connectivity MRI in *Nf1*^+/-^ mice similarly revealed reduced posterior-anterior connectivity along the cingulate cortex as well as disrupted corticostriatal connectivity. The striatum of *Nf1*^+/-^ mice showed increased functional connectivity to somatomotor and frontal cortices and decreased functional connectivity to the auditory cortex. Collectively, these results demonstrate similar alterations across species, suggesting that NF1 pathogenesis is linked to striatal dysfunction and disrupted corticocortical connectivity in the default network.

## Introduction

Neurofibromatosis type 1 (NF1) is an autosomal-dominant tumor pre-disposition syndrome, mostly known for the tendency to develop benign nerve sheath tumors. Beyond the oncological aspect of the disease, hyper-activation of the Ras pathway in NF1 results in several pathologies, both in and outside the CNS (Gutmann et al., 2017). Individuals with NF1, especially at a young age (<21 years), display micro- and macroscopic abnormalities in the central nervous system, and a unique profile of neuropsychological cognitive deficiencies (Hyman et al., 2005; Payne et al., 2010). As attention deficit and learning disabilities are a hallmark of NF1-associated cognitive dysfunction, animal models of NF1 are potentially valuable for clinical research of these phenotypes.

Among the pathology seen in the NF1 brain, are foci of impaired myelination that are associated with cognitive impairment when strategically located (Hyman et al., 2007; Payne et al., 2014). Studies in NF1 genetically engineered mouse models revealed neurotransmitter imbalance and impaired long-term potentiation, which were also reported in humans (Brown et al., 2010; Costa et al., 2002; Cui et al., 2008; Shilyansky et al., 2010). These impairments may underlie NF1-associated behavioral changes that include attention deficits, learning disabilities, hyperactivity, lower IQ, and impaired spatial perception.

The basal ganglia have been implicated in gross phenotypic disruption seen in NF1 pediatric patients and mouse models. Neurofibromin, the protein coded by the *Nf1* gene, has been implicated in regulation of prefrontal and striatal inhibitory networks through increased GABA release (Shilyansky et al., 2010), and motor impairments and structural changes in the striatum measured with MRI have been reported in NF1 mouse models, suggesting a critical role for this region (Petrella et al., 2016; Robinson et al., 2010; van der Vaart et al., 2011). Reduction in striatal dopamine levels has been reported, lending further support for a striatal role in NF1 deficits (Brown et al., 2010). In humans, subtle alterations in structure and function of the basal ganglia were found using histological (Yokota et al., 2008), biochemical (Nicita et al., 2014), and morphological (Duarte et al., 2014) analyses. Functionally, hypo-activation of a frontostriatal circuit was described during working memory related tasks in NF1 patients (Shilyansky et al., 2010). Nevertheless, the impact of neurofibromin deficiency on gross functional organization of brain networks and corticostriatal connectivity remains unclear.

Here we sought to assess the disruption resulting from neurofibromin deficiency using functional imaging, aiming to understand the functional and organizational alterations associated with NF1-specific etiologies. Resting-state functional connectivity MRI (fcMRI) is a non-invasive imaging approach that allows investigation of functional organization of brain networks via coherent spontaneous fluctuations in the blood oxygenation level-dependent (BOLD) fMRI signal. Used in humans (Buckner et al., 2013; Fox and Raichle, 2007) and rodents (Grandjean et al., 2014; Lu et al., 2012; Sforazzini et al., 2014; Stafford et al., 2014; Zerbi et al., 2014) to study brain organization in health and disease, fcMRI enables direct cross-species comparison (Bergmann et al., 2016; Bertero et al., 2018) and provides a path to characterize disease pathogenesis. Leveraging these strengths, we used fcMRI to examine cortical functional organization and corticostriatal connectivity in children with NF1 and in a mouse model of this condition. After we identified abnormal functional connectivity in the pediatric participants, we sought to determine whether these findings are recapitulated in *Nf1*^+/-^ mice using fcMRI in awake passive mice. We found striking similarity of altered corticocortical and corticostriatal functional connectivity recapitulating the disrupted organization found in pediatric participants.

## Materials & Methods

### Ethics statement

The study was approved by the institutional review board of Children’s National Health System. Prior to participation, written assent and consent were obtained from children and their parents/legal guardian respectively, in accordance with the Declaration of Helsinki. All animal experiments were conducted in accordance with the United States Public Health Service’s Policy on Humane Care and Use of Laboratory Animals and approved by the Institutional Animal Care and Use Committee of Technion – Israel Institute of Technology.

### Human Participants

Fourteen children with NF1 (age 9.1 ± 2 years; range 7 – 14; 5 females) were included in this study. All NF1 participants were diagnosed according to the 1987 National Institutes of Health criteria. NF1 participants were scanned at Children’s National Health System on a General Electric whole-body 3T MR scanner (MR750, GE Healthcare, Waukesha, WI). For each participant one or two 6-minute BOLD T_2_^*^ weighted gradient echo-echo planar imaging protocol (GE-EPI; TR/TE 2500/25 ms; Flip angle = 90°; 38 slices, matrix 64 × 64, FOV 240 mm, acquisition voxel size 3.75 × 3.75 × 3.3 mm^3^ ) was acquired, in addition to a high resolution anatomical T1 weighted scan used for registration (TR/TE 9.7/4 ms, 256 × 256 FOV, 160 mm slab thickness, 256 × 256 × 160 matrix at effective resolution of 1 × 1 × 1 mm^3^ , 1 excitation, 12° flip angle). For comparison with healthy subjects, we used healthy participants from the Autism Brain Imaging Data Exchange (ABIDE) database, which is a repository of resting-state fMRI and structural scans of children with or without autism (http://fcon_1000.projects.nitrc.org/indi/abide/). Typically developing children (TDC) from eight centers (NYU, YALE, KKI, STANFORD, UCLA, OLIN, LEUVEN, TRINITY) that were matched to our NF1 group in terms of age and sex were selected. Participants matched for head motion (TDC1) and participants matched for scan parameters (TDC2) were used as comparison groups (see **Table 1**).

**Table 1.** Demographics, movement and scan parameters of the human participants.

|  | NF1 | TDC1 | TDC2 |
| --- | --- | --- | --- |
|  |  | Matched for motion | Matched for scan parameters |
| Age (±SD) | 9.1 (±2.0) | 9.2 (±1.9) | 9.4 (±0.6) |
| N (females) | 14 (5) | 14 (5) | 14 (5) |
| Average Head Displacement (mm) (±SD) | 0.12 (±0.9) | 0.11 (±0.6) | 0.09 (±0.3) |
| TR (s) | 2.5 | 2 | 2.5 |
| Acquisition Resolution (mm<sup>3</sup>) | 3.75×3.75×3.3 | 3×3×3<br>3.4×3.4×4<br>3×3×4<br>3.3×3.3×3.4 | 3.8×3.8×3.8<br>3×3×3 |

### Human Data Preprocessing

Preprocessing followed standard procedures (Yeo et al., 2011) and motion artifacts were minimized using data-scrubbing (Bergmann et al., 2016; Power et al., 2015). Exclusion criteria included framewise displacement exceeding 0.5 mm, temporal derivative RMS variance over voxels (DVARS) exceeding 0.5%, absolute translation over 2 mm or absolute rotation of 0.05°, and exclusion of 1 frame before and two after the detected motion. Five NF1 patients were excluded from this study due to low SNR, excessive movement, or anatomical abnormalities. Whole brain signal, white matter, and ventricle signals were regressed out to minimize signal from non-neural sources, followed by low-pass temporal filtering. Next, data were processed in two parallel pipelines: volume and surface. Volume normalization followed previous procedures (Bergmann et al., 2016) to linearly transform functional images to a downsampled version of the Montreal Neurological Institute (MNI) template at 2 × 2 × 2mm^3^ . Surface normalization followed Yeo et al. (2011) Briefly, structural data was preprocessed using FreeSurfer 4.5.0 (http://surfer.nmr.mgh.harvard.edu). This pipeline constitutes a suite of algorithms that reconstruct a surface representation of the cortical ribbon from an individual participant’s anatomical scan. Group analysis was done through the common spherical coordinate system for both structural and functional data and a 6 mm full width at half-maximum (FWHM) smoothing kernel was applied to the functional data.

### Animal functional imaging procedures

Eleven wild-type (WT; C57BL/6) and eleven NF1 (*Nf1*^+/-^) animals were included. As previously described (Bergmann et al., 2016), mice were implanted with MRI-compatible headposts at age 2-3 months, acclimatized to prolonged head-fixation inside the MRI scanner, and underwent awake head-fixed fcMRI scanning. MRI scans were performed at 9.4 Tesla MRI (Bruker BioSpin GmbH, Ettlingen, Germany) using a quadrature 86 mm transmit-only coil and a 20 mm loop receive-only coil (Bruker). On each scanning session, fast anatomical images were acquired using a rapid acquisition process with relaxation enhancement (RARE) sequence in coronal orientations (30 coronal slices, TR/TE 2300/8.5 ms, RARE factor = 4, flip angle = 180°, 150 × 150 × 450 μm^3^). Whole-brain BOLD functional imaging was achieved using spin echo-echo planar imaging (SE-EPI) sequence (TR/TE 2500/18.398 ms, flip angle = 180°, 30 coronal slices, 150 × 150 × 450 μm; the imaged volume was framed with four saturation slices to avoid wraparound artifacts). Two hundred repetitions were acquired on each run, and four runs were acquired per session. Each animal was scanned over 5–8 sessions (WT: 5.91 ± 0.83; NF1: 5.73 ± 0.467) over a period of 5 to 24 days (WT: 12.27 ± 4.9; NF1: 12.56 ± 4.08; *t*_(20)_ = 0.141, *p* = 0.888). A minimum of zero and a maximum of three scanning sessions were excluded for each animal due to excessive motion (see below for details). A two-tailed unpaired *t*-test did not reveal reliable differences in the number of included sessions per animal (WT: 4.82 ± 0.982; NF1: 5 ± 0.89, *t*_(20)_ = 0.454, *p* = 0.655), number of included frames per animal (WT: 2448.46 ± 657.217; NF: 2350.46 ± 472.636, t(20) = 0.402, *p* = 0.692) or average head motion measured using framewise displacement (WT: 48.3 ± 7.02 μm; NF: 51.4±6.4 μm, *t*_(20)_ = 1.075, *p* = 0.295), indicating that the groups were well-matched.

### Mouse data preprocessing

Raw data were reconstructed using ParaVision 5.1 (Bruker). Standard mouse fMRI preprocessing was based on previous work (Bergmann et al., 2016; Kahn et al., 2011), and included removal of the first two volumes for T_1_-equilibration effects, compensation of slice-dependent time shifts, rigid body correction for motion within and across runs, registration to a downsampled version (150 × 150 × 450 μm^3^ ) of the Allen Mouse Brain Connectivity (AMBC) Atlas which includes anatomical annotations (Lein et al., 2007) and optical density maps from anatomical tracing experiments (Oh et al., 2014), and intensity normalization.

Preprocessed data were then passed for additional fcMRI-specific processing steps. A data scrubbing protocol was applied to eliminate motion-related artifacts (Bergmann et al., 2016; Power et al., 2015). Exclusion criteria framewise displacement of 50 μm and DVARS of 150% inter-quartile range (IQR) above the 75^th^ percentile, with exclusion of 1 additional frame after the detected motion. Runs with fewer than 50 frames and sessions with fewer than 300 frames were excluded. A total of 20 out of 128 sessions were excluded (WT = 12/65; NF1 = 8/63). Next, data underwent demeaning, detrending and regression of sources of spurious or regionally nonspecific variance including six motion parameters, average time courses in the ventricles and global signal, with temporally shifted versions of these waveforms removed by inclusion of their first temporal derivatives. Temporal filtering was used to retain frequencies between 0.009–0.08 Hz and data were spatially smoothed using a 600 μm FWHM Gaussian blur.

### Experimental design and statistical analysis

#### Human data analysis

For each group of participants, we used a global similarity-based clustering approach to parcellate the cortex into seven networks (Yeo et al., 2011). The Sørensen–Dice index (also termed *similarity coefficient*), calculated as *D* = 2 × (A ∩ B) / (|A| + |B|), was used to quantify the overlap between NF1 (A) and TDC (B) cortical parcellation, where a similarity coefficient of 1 (*D*) represents a perfect overlap. A complementary seed-based analysis was conducted using a ~2 cm^3^ sphere located in the posterior cingulate cortex in which seed-based Fisher’s Z transformed *r* correlation map submitted to a one-way *t*-tests (SPM, Wellcome Department of Cognitive Neurology, London, UK). In addition, seed-to-seed analysis examined the connectivity between the posterior and anterior cingulate cortex. A similar seed-to-seed approach was used to characterize corticostriatal connectivity by calculating the Z transformed *r* correlation between striatal seeds (in volume space) and cortical seeds (in surface space), as previously described (Choi et al., 2012).

#### Mouse data analysis

To estimate functional connectivity in the mouse brain, 450 μm-diameter spheres (5 voxels) were placed at the center of the relevant AMBC tracing experiments to define regions-of-interest (ROIs), as previously described (Bergmann et al., 2016). In each mouse, seed-based Fisher’s Z transformed *r* correlation maps were averaged across sessions, and the averaged maps were submitted to a one-way *t*-test (SPM). These maps were used to characterize posterior-anterior connectivity in the cingulate cortex. In addition, a seed-to-seed analysis was used to formally test changes in posterior-anterior connectivity along the cingulate cortex and to characterize corticostriatal connectivity. Striatal seeds were defined based on the anatomical connectivity profile from the AMBC Atlas, as previously described (Grandjean et al., 2017; Oh et al., 2014).

## Results

We first explored the impact of neurofibromin deficiency on brain-wide functional organization in pediatric participants. For this we utilized a dataset of children with NF1 and two age- and gender-matched comparison groups of typically developing children (TDC). We addressed two potential confounds that might mask veridical differences linked to NF1: participant head motion (TDC1) and scan parameters (TDC2).

### Organization of brain-wide functional connectivity in NF1 children

To examine the brain-wide organization, we utilized a global similarity-based clustering parcellation. In both comparison groups parcellation to seven networks yielded Default, Visual, Frontoparietal, Limbic and Dorsal Attention (DA) networks that were similar to those in adults (Yeo et al., 2011). The Somatomotor (SM) and Ventral Attention (VA) parcellations differed relative to adults, not showing the formation of the typical interdigitated, distributed areas that form the VA and the contiguous post- and pre-central sulcus parcel that is observed in adults (**Fig. 1A**). In the NF1 group, the same, immature, parcellation pattern of the SM and VA was evident, as well as the typical formation of the visual network. However, the NF1 group failed to demonstrate the proper organization of other association networks including Default, Frontoparietal, DA, and Limbic (**Fig. 1A**). Quantitative comparison of network overlap between the two control groups revealed close agreement with a median Dice index of 0.87 (range 0.75–0.96) in both the right and left hemispheres (**Fig. 1B**). Examining network overlap between NF1 and TDC1 revealed lower similarity coefficients for Default, Frontoparietal, DA, and Limbic networks, all networks that are usually characterized by patterns of distal connectivity, while VA, SM and Visual networks had similarity coefficients greater than 0.8 (**Fig. 1B**). In these latter networks, distal connectivity is weaker relative to local connectivity (Sepulcre et al., 2010). Further, examination of the spatial extent of Default and Frontoparietal networks revealed that while their parcellations differs, when considered together they demonstrate fair overlap (**Fig. 1C**). Network overlap between NF1 and TDC2 demonstrated similar results with decreased overlap of distributed association networks (Default: 0.48, Frontoparietal: 0.48, DA: 0.33, and Limbic: 0.55), while other, less dispersed networks demonstrated higher overlap (SM: 0.79, VA: 0.82, and Visual: 0.89). Together, these findings indicate disrupted functional connectivity in association networks in NF1 children.

**Figure 1.**
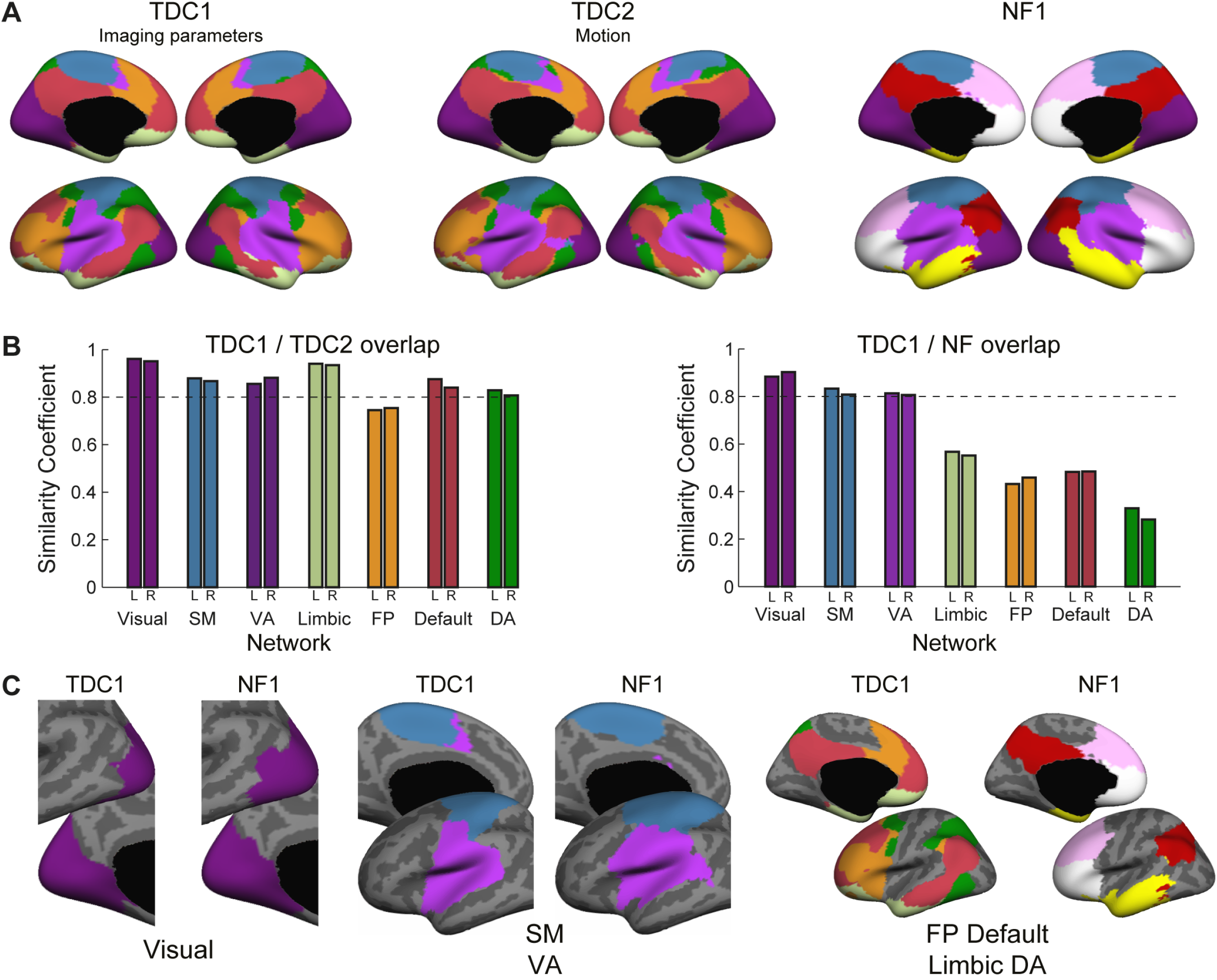
Pediatric NF1 participants demonstrate disrupted association network organization. (A) A global similarity-based clustering parcellation to seven networks of two age- and gender-matched typically developing children (TDC) comparison groups, TDC1 group (*left*) matched for MRI repetition time and TDC2 group (*middle*) matched for participant motion. Network parcellation of the NF1 group (*right*) demonstrates a notable lack of long-distance connectivity. (B) Network overlap estimated with Sørensen–Dice index (similarity coefficient) between the two comparison groups (all greater than 80%; *left*) reveals close agreement between left and right hemispheres. The similarity coefficient between TDC1 and NF1 groups (*right*) is virtually equivalent for the Visual, Ventral Attention (VA) and Somatomotor (SM), but low for Limbic, Dorsal Attention (DA), Frontoparietal (FP) and Default networks. (C) Consistent with the similarity coefficient, spatial extent is conserved for the Visual network and is similar to the adult form (not shown; cf. Yeo et al. 2011). Putative SM and VA networks are similar across TDC1 and NF1 but differ relative to adults. The DA, FP, Default and Limbic networks differ between TDC1 and NF1 although the spatial extent of the two networks is similar when considered together.

To better characterize the Default network, we used a seed-based functional connectivity analysis in the posterior cingulate cortex (**Fig. 2A**). The results confirm the global similarity clustering findings with the NF1 group showing reduced posterior-anterior connectivity along the cingulate cortex. To formally test whether the reduced connectivity is associated with physical distance, we placed four seeds along the anterior cingulate cortex, and calculated their correlation with the posterior cingulate seed (**Fig. 2B**). Repeated-measures ANOVA revealed a significant main effect of Group (F_(1,76)_=4.71, p = 0.039) as well as a significant interaction between Group and Longitudinal Axis (F_(3,78)_ = 3.72, *p* =0.028, ε_H-F_ = 0.717), confirming distance-dependent reduced connectivity in the NF1 group.

**Figure 2.**
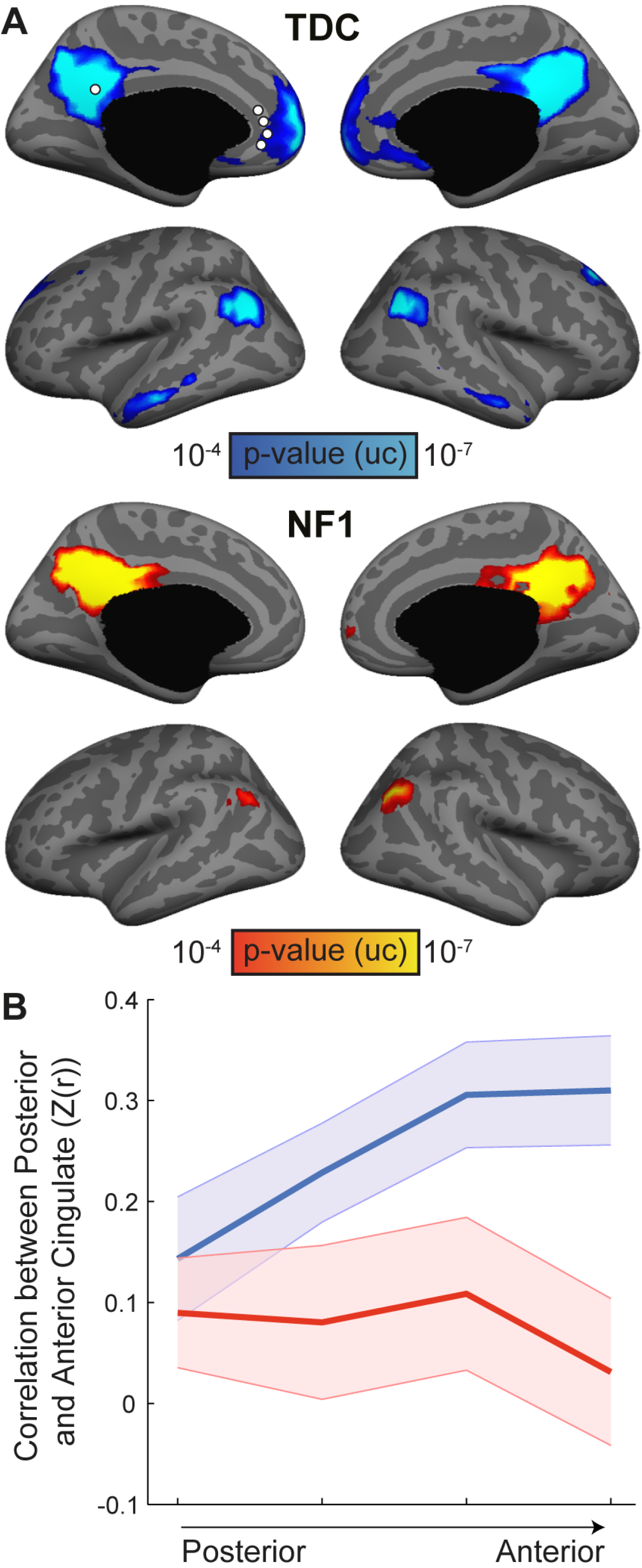
Reduced posterior-anterior connectivity along the cingulate cortex in NF1 children. (A) Seed-based analysis for the posterior cingulate cortex in TDC (*left*) and NF1 (*right*) demonstrates reduced long range functional connectivity in the TDC group. (B) Connectivity profile between the posterior cingulate cortex and a series of seeds along the anterior cingulate cortex revealed significant main effect of Group and an interaction between Group and Longitudinal axis of the cingulate cortex, indicating reduced distance-dependent connectivity in NF1 children.

### Corticostriatal connectivity and topographic striatal organization are altered in NF1

Since the most dramatic corticocortical changes were observed in association networks, we focused our investigation of corticostriatal connectivity in these cortical systems. To investigate corticostriatal connectivity patterns, we replicated a seed-to-seed analysis from Choi et al. (2012). The analysis is based on 16 cortical seeds located in different association systems and three striatal seeds that are known to be connected to the Default, Frontoparietal, and Limbic networks (**Fig. 3A**). Examining the connectivity profile of each of the striatal seeds (**Fig. 3B**), we found that overall, the TDC1 group shows similar patterns to the original analysis in adults (Choi et al., 2012, cf. Fig. 12), with some differences in the Default region of the striatum which failed to demonstrate the mature connectivity profile. Comparing the results to the NF1 group, we found that NF1 children show decreased corticostriatal connectivity in the cortical regions of the Frontoparietal (PFCmp, PFCla, PGa) and Default (PFCdp, PFCm, PGc, PCC) networks, as well as increased connectivity with the Limbic subgenual cingulate cortex (scg25). To formally test the topography of corticostriatal connections and to compare between groups we focused on the Default, Frontoparietal and Limbic cortices, and averaged the corticostriatal connectivity of all seeds that belong to the same network based on the definition in adults (**Fig. 3C**); then, we submitted the results to a repeated-measure ANOVA. First, we found a significant interaction between Striatal Seed and Cortical Network (F_(4,104)_ = 21.09, *p* < 0.001, ε_H-F_ = 0.745), confirming topographical organization of different cortical networks within the striatum. Next, we found a significant interaction between Cortical Network and Group (F_(2,52)_ = 6.67, *p* = 0.008, ε_H-F_ = 0.764) but not a three-way interaction between Striatal Seed, Cortical Network and Group (F_(4,104)_ = 0.94, *p* = 0.43, ε_H-F_ = 0.745), indicating alterations that do not change to topography of corticostriatal connectivity. To break down the interaction between Striatal Seed and Cortical Networks, we conducted similar analysis for each pair of networks using Bonferroni adjusted alpha levels of .0133 per test (.05/3). We found significant interaction when comparing Limbic and Frontoparietal networks (F_(1,26)_ = 8.59, *p* = 0.0077, ε_H-F_ = 1), with no interactions between these and the Default network (Default vs. Limbic: F_(1,26)_ = 5.74, *p* = 0.024, ε_H-F_ = 1; Default vs. Frontoparietal: F_(1,26)_ = 2.2, *p* = 0.15, ε_H-F_ = 1). Collectively, these results indicate increased limbic and decreased frontoparietal corticostriatal connectivity in NF1 pediatric patients.

**Figure 3.**
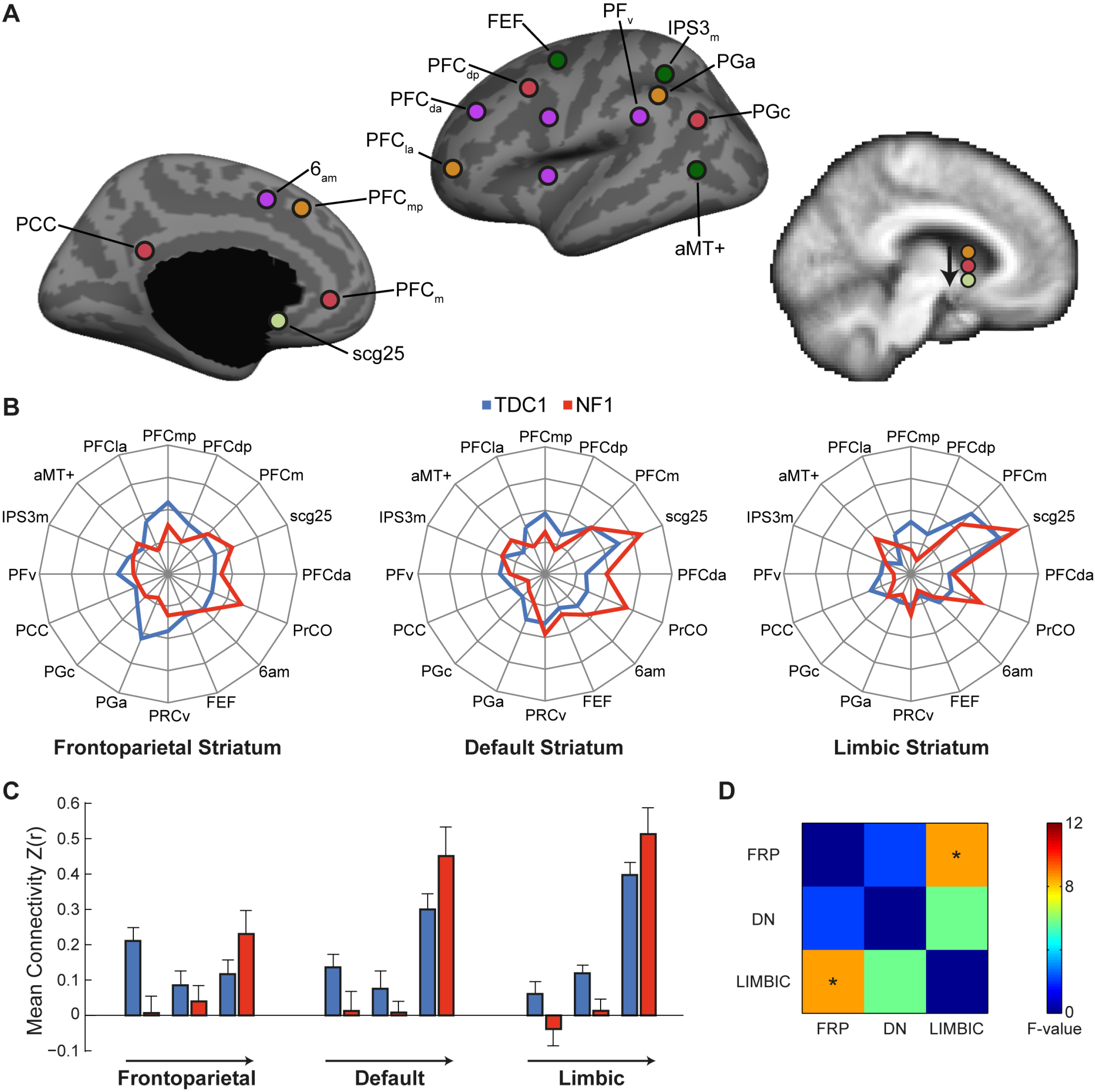
Corticostriatal functional connectivity in NF1 and typically developing children (TDC). (A) Seed locations in the human cortex and striatum. (B) Three polar plots display corticostriatal functional connectivity of the three striatal sub-regions (frontoparietal, default and limbic). Polar scales ranges from *z*(*r*) of −0.2 to −0.6 in 0.2 increments. (C) Bar plots depict average corticostrital connectivity between each cortical network and the different striatal seeds. Error bars indicate standard errors of the mean. (D) Quantification of the interaction between pairs of Cortical Network and Striatal Seed, \**p* < 0.05, corrected for multiple comparisons using the Bonferroni correction.

### NF1 mice demonstrate altered posterior–anterior connectivity along the cingulate cortex

Next, we examined whether the altered functional organization we found in NF1 pediatric participants is recapitulated in an NF1 mouse model. Since the global similarity clustering algorithm has yet to bet adapted to mice, we focused on the disrupted posterior-anterior connectivity, which was previously reported in rodents, and considered homologous (Lu et al., 2012; Stafford et al., 2014). Seed-based functional connectivity analysis of the retrosplenial cortex showed that NF1 mice fail to demonstrate posterior-anterior connectivity (**Fig. 4A**). To formally test this observation, we used seed-to-seed analysis between the retrosplenial cortex and four seeds along the anterior cingulate area (**Fig. 4C**). The connectivity profiles were submitted to a repeated-measures ANOVA, which revealed a significant main effect of Group (F_(1,20)_=6.1, *p* = 0.023) as well as a significant interaction between Group and Longitudinal Axis (F_(3,60)_ = 3.53, *p* =0.036, ε_H-F_ = 0.711) indicating distance-dependent reduced connectivity in the NF1 group, recapitulating the human phenotype.

**Figure 4.**
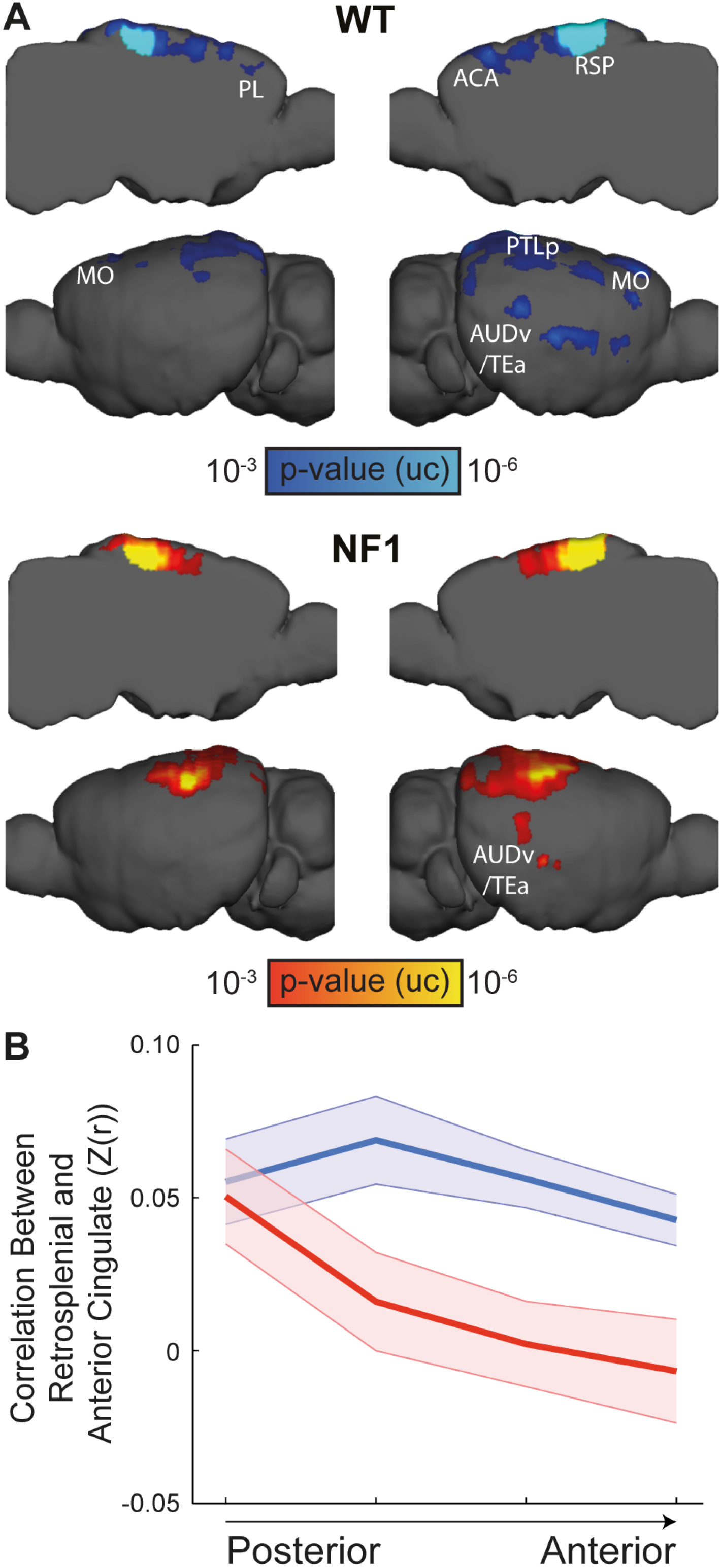
Reduced posterior-anterior connectivity along the cingulate cortex in NF1 mice. (A) Seed-based analysis for the retrosplenial cortex in WT (*left*) and NF1 (*right*) mice demonstrates posterior-anterior functional connectivity only in the WT group. (B) Connectivity profile between the retrosplenial cortex and a series of seeds along the anterior cingulate area revealed significant main effect of Group and an interaction between Group and Longitudinal axis of the cingulate cortex, indicating reduced distance-dependent connectivity in NF1 children. ACA – anterior cingulate cortex; AUDv/Tea – ventral auditory/temporal association cortices; MO – motor cortex; PL – prelimbic cortex; PTLp – posterior parietal association cortex; RSP – retrosplenial cortex.

### NF1 mice demonstrate disrupted cortcostriatal connectivity

After we found similar alteration in corticocortical connectivity, we sought to examine whether NF1 mice recapitulate the disrupted cortcostriatal connectivity we found in humans. Unlike the mammalian cerebral cortex, which underwent dramatic expansion and elaboration (Geschwind and Rakic, 2013), the striatum has remained relatively conserved (Grillner et al., 2013), allowing a more straightforward comparison across species. Therefore, we hypothesized that NF1 mice will demonstrated altered organization similar to that seen in humans. A recent study (Grandjean et al., 2017) investigated corticostriatal connectivity in the mouse brain by comparing fcMRI data to anatomical tracing data from the AMBC Atlas. This study reported a close agreement between anatomical and functional corticostriatal connectivity, as both modalities demonstrated a similar topographical organization of auditory, somatomotor and frontal cortices along the longitudinal axis of the mouse striatum. Therefore, we took a similar approach and used the AMBC Atlas to guide our seed-based analysis.

Based on the reorganization of the human NF1 striatum, we hypothesized that corticostriatal functional connectivity in NF1 mice would be altered. To test this hypothesis, we placed seeds in 16 cortical regions. Representative anatomical and functional maps for the ventral auditory, secondary somatosensory, primary motor and anterior cingulate cortices demonstrated close agreement between anatomical and functional connections in both WT and NF1 mice (**Fig. 5**), as functional connectivity maps generally captured inter-hemispheric corticocortical and both inter- and intra-hemispheric corticostriatal functional connectivity.

**Figure 5.**
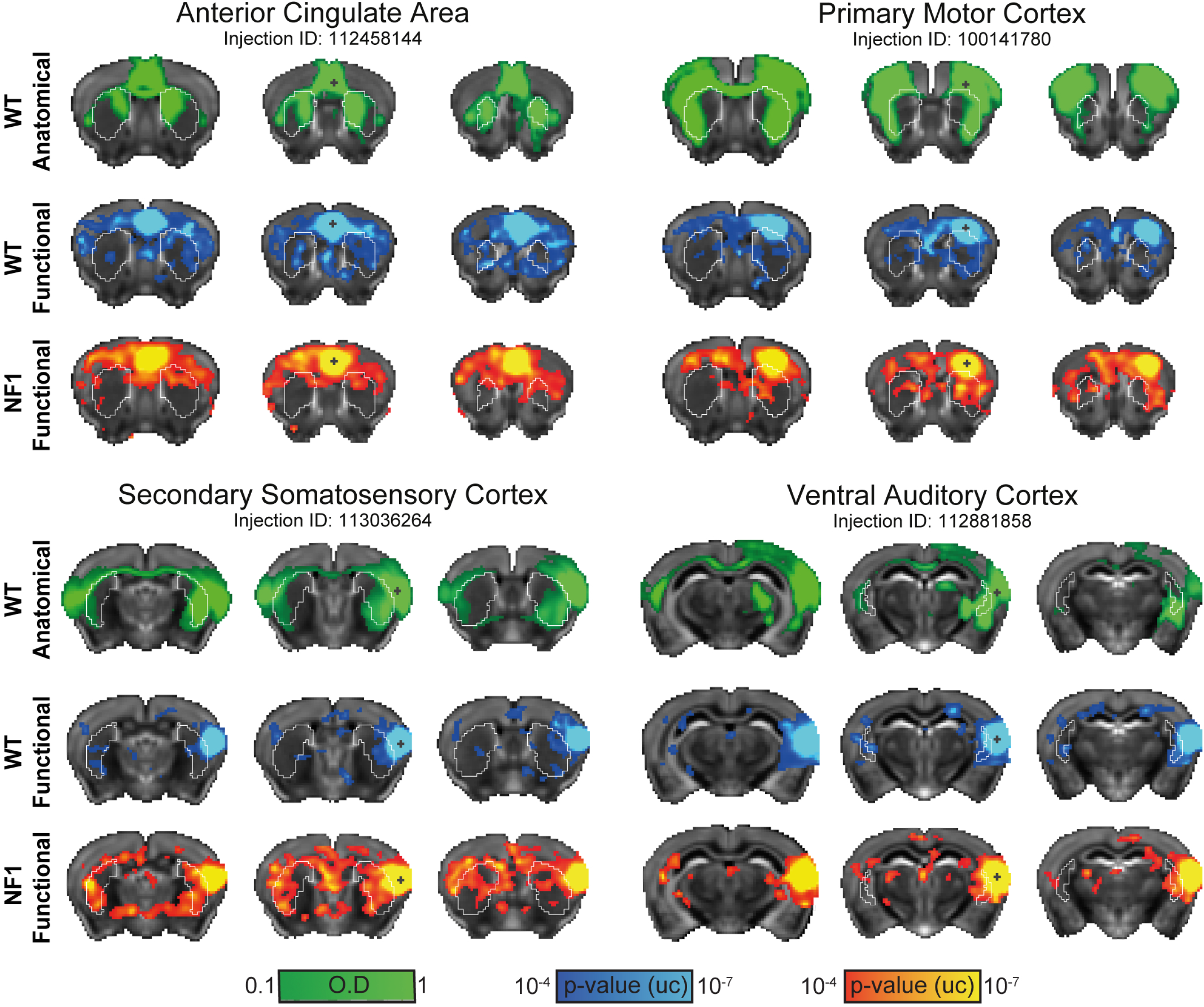
Functional and anatomical corticostriatal connectivity in the mouse brain. Qualitative comparisons between optical density maps (green) representing anatomical connectivity, and statistical parametric maps (*p* < 0.0001, voxel extent ≥ 5) representing positive functional connectivity correlations in WT (blue-light blue) and NF1 (red-yellow) mice, are presented for four cortical regions. Overlapping corticocortical and corticostriatal connections can be observed in WT but to a lesser extent in NF1. The parametric maps are overlaid on a downsampled version of the Allen Mouse Brain Atlas that matches at the original fMRI resolution.

To better characterize the organization of corticostriatal functional connectivity in WT and NF1 mice, we used the anatomical connectivity data from the AMBC Atlas to define seeds in three striatal subregions (**Fig 6A**), which are known to be connected to frontal, somatomotor and auditory cortices, respectively. We assigned our 16 ROIs to five cortical systems: frontal, somatomotor, auditory, parietal association, and visual (**Fig. 6A**), and calculated the connectivity profiles of the different striatal subregions (**Fig. 6B**). We found that overall the WT group show distinct topography with different cortical seeds peaks at different striatal subregions, as previously reported (Grandjean et al., 2017). Comparing the results to the NF1 group, we found that NF1 mice show decreased corticostriatal connectivity in the auditory cortices (AUDd, AUDp, AUDv) and increased connectivity with the prefrontal cortices (ACA, ORBm, PL), as well as some somatomotor cortices (SSp-bfd, SSs). To formally test the topography of corticostriatal connections and to compare between groups we focused on frontal, somatomotor and auditory cortices, and averaged the corticostriatal connectivity of all seeds that belong to the same network (**Fig. 6C**); then, we submitted the results to a repeated-measure ANOVA. First, we found a significant interaction between Striatal Seed and Cortical Network (F_(4,80)_ = 41.94, *p* < 0.001, ε_H-F_ = 1), confirming topographical organization of different cortical networks within the striatum. Next, we found a significant interaction between Cortical Network and Group (F_(2,40)_ = 12.74, *p* = < 0.001, ε_H-F_ = 0.816) and three-way interaction between Striatal Seed, Cortical Network and Group (F_(4,80)_ = 3.17, *p* = 0.018, ε_H-F_ = 1), indicating topographical organization of corticostriatal organization which is altered in NF1 mice. To break down the interaction between Striatal Seed and Cortical Networks, we conducted similar analysis for each pair of networks using Bonferroni adjusted alpha levels of .0133 per test (.05/3). We found significant interaction between Auditory and both Frontal (F_(1,20)_ = 26.56, *p* < 0.001, ε_H-F_ = 1) and Somatomotor (F_(1,20)_ = 11.19, *p* = 0.003, ε_H-F_ = 1) networks, with no interactions between these and the latter (Frontal vs. Somatomotor: F_(1,20)_ = 0.12, *p* = 0.737, ε_H-F_ = 1). Collectively, these results indicate decreased auditory and increased somatomotor and frontal corticostriatal connectivity in NF1 mice, demonstrating similar phenotype across species.

**Figure 6.**
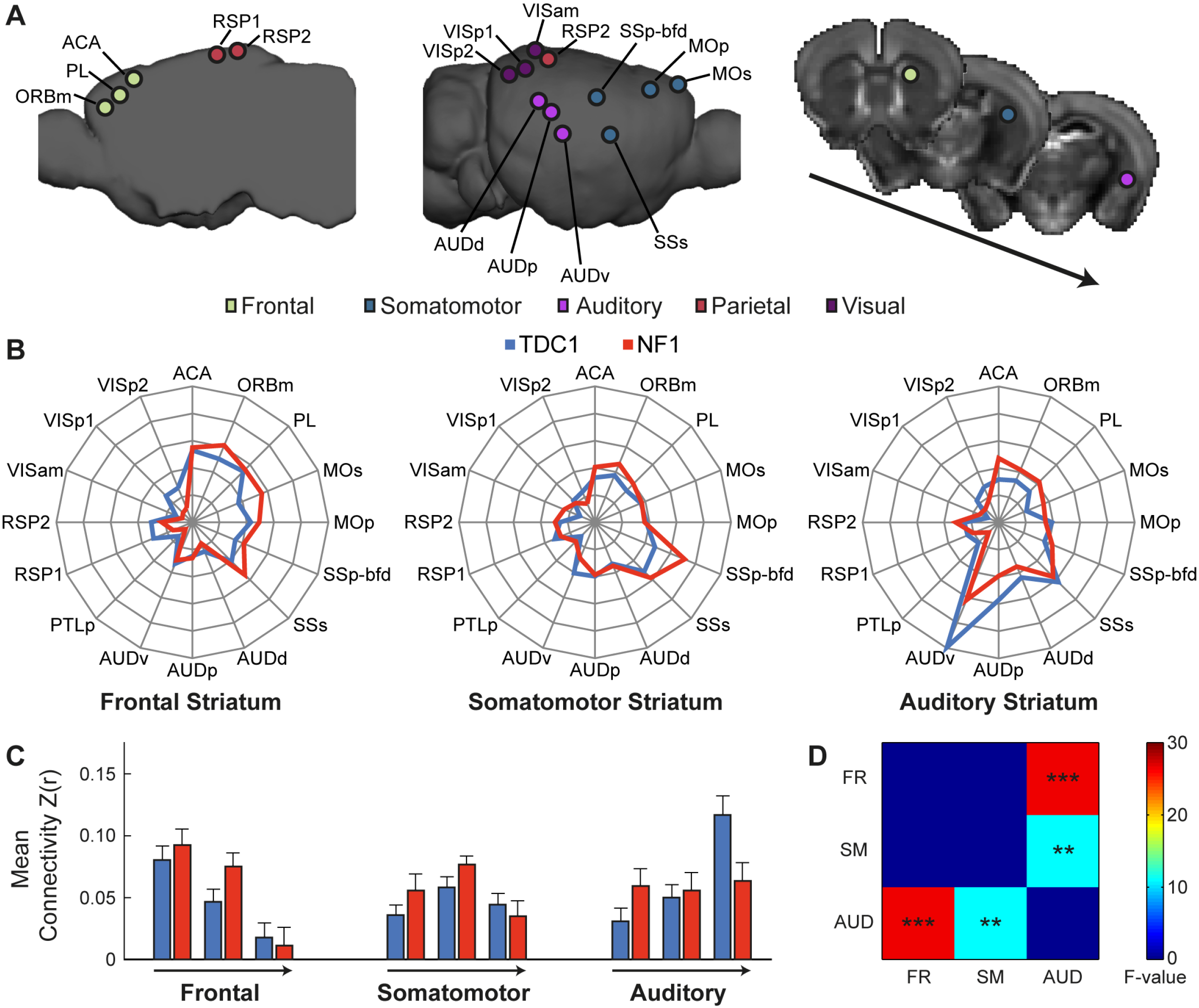
Alterartion of corticostriatal functional connectivity in NF1 mice. (A) Seed locations in the mouse cortex and striatum. (B) Three polar plots display corticostriatal functional connectivity of the three striatal subregions (frontal, somatomotor and auditory). Polar scales range from *z*(*r*) of −0.05 to −0.2 in 0.05 increments. (C) Bar plots depict average corticostrital connectivity between each cortical network and the different striatal seeds. Error bars indicate standard errors of the mean (D) Quantification of the interaction between pairs of Cortical Network and Striatal Seed (frontal [FR], somatomotor [SM] and auditory [AUD]), \*\**p* < 0.01, \*\*\**p* < 0.001, corrected for multiple comparisons using the Bonferroni correction. AUD – auditory cortex: dorsal (d), primary (p), ventral (v); MO – motor cortex: primary (p), secondary (s); ORBm – medial orbitofrontal cortex; PL – prelimbic cortex; PTLp – posterior parietal association cortex; RSP – retrosplenial cortex; SSp-bfd – barrel-related primary somatosensory cortex; SSs – secondary somatosensory cortex; VIS – visual cortex: anteromedial (am), primary (p).

## Discussion

Here, we report a cross-species comparative analysis of the functional organization of the NF1 brain in both humans and mice. In typically developing children, we characterized the organization of large-scale cortical functional networks and formation of corticostriatal functional connectivity in this age group for the first time, and identified alterations in functional organization associated with NF1. We then applied the same functional imaging method in a mouse model of NF1 and found similar alterations in corticocortical and corticostriatal connectivity.

Hyper-activation of the Ras pathway results in several pathophysiological alterations that may lead to the typical NF1 cognitive phenotype. We discovered two fundamental disruptions that may be explained by cellular defects known to result from neurofibromin deficiency. First, we identified that the development of large-scale association networks, the default and frontoparietal, differs in NF1 relative to healthy participants, and that this pattern is associated more broadly with disrupted antero-posterior functional connectivity in the cingulate cortex. This disruption was replicated in the mouse data, and may be explained by aberrant myelination known to occur in both NF1 mouse models and pediatric patients. Second, we demonstrated altered organization of corticostriatal functional connectivity in humans and mice, as both species showed differential modulations of distinct cortical networks. This is among few demonstrations of a common pathological functional organization in humans and a genetic mouse model of the human condition. These data highlight the importance of neurofibromin signaling in the establishment of cortical association and corticostriatal networks.

Analysis of cortical functional organization in typically developing children revealed nascent immature organization of the somatomotor and ventral attention cortical networks. In adults the somatomotor network covers the entire primary motor and somatosensory cortices, extends ventrally to the insular area, and includes parts of the auditory cortex, while the ventral attention network is dispersed, and includes the inferior frontal gyrus, supramarginal gyrus, superior temporal gyrus (posterior part), and mesially the cingulate gyrus (Yeo et al., 2011). Here, typically developing children demonstrated conserved division into networks when these were estimated using the same global similarity-based clustering parcellation (Yeo et al., 2011) with the exception that the somatomotor and ventral attention networks were divided into dorsal and ventral components. This result was similar in both comparison groups with high inter-group overlap of both components. This finding of immature organization of the somatomotor and ventral attention networks in healthy children is compatible with current concepts of the formation of distinct, dispersed, functional connectivity networks with age. Stevens et al. (2009) analyzed a cohort of 100 participants aged 13-30 and reported a decrease in between-network functional connectivity and an increase in within-network functional connectivity that correlated with age, consistent with current descriptions of age-related differences (Dennis and Thompson, 2013). This nascent parcellation of networks was recapitulated in the NF1 group, demonstrating age-appropriate organization of these networks, as well as the mature organization of the visual and limbic networks in both healthy and NF1 participants.

Long-range functional connectivity and distributed functional networks are known to develop slowly in parallel with the formation of myelin from childhood to adulthood (Supekar et al., 2009). Impairment in the typical maturation process was recently suggested in NF1. Tomson et al. (2015) reported impairment in antero-posterior functional connectivity and suggested that it was related to malformation of association networks in NF1 adults. This finding was replicated in our study in a cohort of NF1 pediatric patients in two independent analyses. First, unlike in TDC, the NF1 group failed to demonstrate the formation of age-appropriate association networks that are based on antero-posterior functional connectivity (default, frontoparietal and dorsal attention). Further, a seed-based cortical functional connectivity analysis demonstrated weaker correlations of default network regions in NF1. The latter analysis was replicated in NF1 mice, which also demonstrated disrupted posterior-anterior connectivity along the cingulate cortex. This disruption in the formation of long-range connectivity may be related to disruptions in myelination, which is essential for structural maturation of the human brain (Dubois et al., 2008), since the *Nf1* gene has been implicated in white matter defects including myelin decompaction (Lopez-Juarez et al., 2017; Mayes et al., 2013; Rosenbaum et al., 1999). In direct relation to these microscopic alterations, studies in human NF1 patients demonstrated broadly abnormal myelination, particularly in the frontal lobe, expressed as significantly decreased radial diffusivity measured with diffusion weighted imaging (Karlsgodt et al., 2012). These findings suggest that the reduced long-range functional connectivity of the dorsal attention, frontoparietal and default networks observed here might be related to disruption in myelination. Moreover, our results are in close agreement with other studies that linked reduced posterior-anterior connectivity in the Default network and neurodevelopmental disorders in humans and mice (Bertero et al., 2018; Liska et al., 2018; Pagani et al., 2018; Washington et al., 2014; Yerys et al., 2015).

The striatum has been implicated in the NF1 cognitive phenotype in NF1 mouse models and human patients (Brown et al., 2010; Nicita et al., 2014; Petrella et al., 2016; Yokota et al., 2008). We sought to characterize whether corticostriatal functional connectivity was altered in NF1. An in-depth mapping of striatal functional connectivity in TDC relative to adults revealed that maturation was correlated with a developmental shift from preferential functional connectivity of the basal ganglia and the somatomotor network in childhood, to the frontoparietal and ventral attention association networks in adulthood (Greene et al., 2014). In our study, while typically developing children displayed mature and balanced connectivity profiles in limbic and frontoparietal striatal regions, the NF1 group showed increased limbic and decreased frontoparietal connectivity. This finding is consistent with a previous work (Shilyansky et al., 2010) that implicated prefrontal and striatal functional connectivity in working memory impairment in adults with NF1.

Similar to our findings in humans, *Nf1*^+/-^ mice demonstrated an overall conserved topographic functional organization frontal and somatomotor striatal region, with markedly altered topography in the auditory striatum. In addition, NF1 mice showed increased somatomotor and frontal with decreased auditory corticostriatal connectivity. The fact that both children and mice show differential alterations in corticostriatal connectivity suggests that mesoscopic phenotype found in both species may be used to dissect the mechanisms governing these disruptions. This result suggests that therapies that reduce increased corticostriatal functional connectivity may be beneficial in alleviating the NF1 cognitive phenotype. Further, longitudinal pre-clinical functional imaging, which is possible in awake mouse fMRI (Bergmann et al., 2016), will complement demonstrations of mapping (Lu et al., 2012; Stafford et al., 2014) and changes in brain-wide functional connectivity in models of disease (Bertero et al., 2018; Grandjean et al., 2014; Zerbi et al., 2014).

The human striatum demonstrates a consistent, topographically arranged, functional connectivity pattern (Choi et al., 2012; Jaspers et al., 2017). Alterations in the functional organization of the basal ganglia were reported in both ASD and ADHD. A study comparing ASD patients with a typically developing matched cohort found that the ASD group failed to demonstrate the age associated reduction of functional connectivity between the putamen and the frontal pole (Balsters et al., 2017). In a study focusing on adolescents with ADHD, increased correlation of the putamen and ventral striatum with several cortical areas including motor and prefrontal cortices was shown (Oldehinkel et al., 2016). Studies of mouse models of ASD reported altered corticostriatal connectivity (Peixoto et al., 2016; Wang et al., 2016; Zerbi et al., 2018), and in a study on the 16p11.2 microdeletion model of autism, impaired prefrontal functional connectivity was demonstrated (Bertero et al., 2018). As many as 40% of NF1 patients display ASD symptomatology (Walsh et al., 2013), and 60-70% display the ADHD phenotype (Coude et al., 2007). Our findings of altered corticostriatal and decreased long-range corticocortical functional connectivity, with a possible key role for myelin dysfunction (Lopez-Juarez et al., 2017; Mayes et al., 2013; Rosenbaum et al., 1999), suggest a potential common disruption pathway pivotal for this phenotype in both NF1 and ASD. These findings warrant further investigation at the individual participant level by clinically and dimensionally assessing ASD and ADHD phenotypes in NF1 patients.

## Acknowledgements

This work was supported by the Israel Science Foundation (770/17; to I.K.), the National Institutes of Health (1R01NS091037; to I.K.), the Adelis Foundation (to I.K.) and Gilbert Family Neurofibromatosis Foundation (to R.J.P.). We thank Nancy Ratner for helpful comments on an early version of the manuscript, Technion’s Biological Core Facilities and Edith Suss-Toby for her assistance with MRI, and the Technion Preclinical Research Authority, and Nadav Cohen for assistance with animal care.

## Author contributions

B.S., E.B., S.C., and I.K. designed research; B.S. E.B., G.Z., J.A, A.K., R.J.P., L.G.V., M.T.A. performed research; B.S., E.B., G.Z., and I.K. analyzed data; and B.S., E.B., G.Z., N.B., F.X.C., L.B.-S., R.J.P, L.G.V., S.C., M.T.A., and I.K. wrote the paper.

